# Increasing the gradient of energetic cost does not initiate adaptation in human walking

**DOI:** 10.1101/2020.05.20.107250

**Authors:** Surabhi N. Simha, Jeremy D. Wong, Jessica C. Selinger, Sabrina J. Abram, J. Maxwell Donelan

## Abstract

When in a new situation, the nervous system may benefit from adapting its control policy. In determining whether or not to initiate this adaptation, the nervous system may rely on some features of the new situation. Here we tested whether one such feature is salient cost savings. We changed cost saliency by manipulating the gradient of participants’ energetic cost landscape during walking. We hypothesized that steeper gradients would cause participants to spontaneously adapt their step frequency to lower costs. To manipulate the gradient, a mechatronic system applied controlled fore-aft forces to the waist of participants as a function of their step frequency as they walked on a treadmill. These forces increased the energetic cost of walking at high step frequencies and reduced it at low step frequencies. We successfully created three cost landscapes of increasing gradients, where the natural variability in participants’ step frequency provided cost changes of 3.6% (shallow), 7.2% (intermediate) and 10.2% (steep). Participants did not spontaneously initiate adaptation in response to any of the gradients. Using metronome-guided walking— a previously established protocol for eliciting initiation of adaptation—participants next experienced a step frequency with a lower cost. Participants then adapted by −1.41±0.81 (p=0.007) normalized units away from their originally preferred step frequency obtaining cost savings of 4.80±3.12%. That participants would adapt under some conditions, but not in response to steeper cost gradients, suggests that the nervous system does not solely rely on the gradient of energetic cost to initiate adaptation in novel situations.

## Introduction

We routinely perform movements in a variety of situations. This includes handling of different-sized objects, walking on uneven terrain, or running with fatiguing muscles. Some of these situations are familiar, and for these situations, our nervous system may have already learned an optimal, or near-optimal, control policy (Izawa et al., 2008; Wolpert et al., 2011; Wolpert and Flanagan, 2016). In the task of walking on a treadmill, for example, people can rapidly select the step frequency that minimizes energetic cost for each new walking speed (Pagliara et al., 2014; Snaterse et al., 2011). But in novel situations, the nervous system hasn’t had the experience to determine whether an existing policy remains optimal, or if a new policy would be better (Wolpert et al., 2011; Wolpert and Flanagan, 2016). To determine this, the nervous system must adapt the existing policy and experience the outcome (Sutton et al., 1992; Wolpert et al., 2011). This adaptation is beneficial only when there is a new optimal solution, the presence of which the nervous system does not know in advance. If the old policy remains the optimal policy, then the act of adapting to new policies is itself sub-optimal—the nervous system would benefit most by exploiting its existing control policy (Sutton et al., 2017). In this paper, we aim to identify a feature of novel situations that cues the human nervous system to initiate adaptation of its control policy.

Our nervous systems do not always initiate adaptation in novel situations. In reaching experiments, people typically initiate adaptation when presented with a force-field that creates a novel relationship between cost and control policy (Shadmehr and Mussa-Lvaldi, 1994; Wolpert et al., 2011). However, when this is followed by another force-field that creates a different novel relationship, the nervous system reverts to erroneously exploiting its original control policy (Gupta and Ashe, 2007; Wolpert et al., 2011). Similar interference to adaptation is also observed in studies that create novel situations using visuomotor rotations or reversals (Krakauer et al., 2019). In walking tasks, exoskeletons designed to improve walking economy can underperform partly because people are unable to adapt their gait to take full advantage of the benefits that the exoskeleton can offer (Jackson and Collins, 2015; Wong et al., 2019; Zhang et al., 2017). In split-belt walking, people do not adapt their step lengths back to baseline when the speeds of the two belts are changed gradually (Roemmich and Bastian, 2015). However, in all of these tasks, the nervous system can and does adapt when certain modifications are made to the novel situations (Krakauer et al., 2019; Selinger et al., 2015; Torres-Oviedo et al., 2011; Wolpert and Flanagan, 2016; Zhang et al., 2017). This suggests that the nervous system relies on particular features of the novel situations to determine if and when to initiate adaptation.

One potential feature used by the nervous system to initiate adaptation is salient cost savings. Here we use *cost savings* to refer to an improvement in the nervous system’s objective function. This may be decreased energetic cost, increased stability, increased accuracy, or some combination of these and other contributors to the objective function. *Saliency* refers to how clear it is to the nervous system that cost savings can be gained, and how it should adapt its control policy to gain the savings. As illustrated in Figure 1, saliency depends on at least three factors. First, execution variability about the nominal policy—due to either imperfect execution, purposeful exploration, or guidance by an external input— allows the nervous system to experience a greater range of cost savings if they exist (Figure 1B). Second, measurement noise decreases the ability of the nervous system to discern the presence of cost savings (Figure 1C). Third, for any given execution variability and measurement noise, an increase in the gradient of the cost landscape increases the ability of the nervous system to discern a cost savings (Figure 1D). If cost savings are not salient— be it due to any combination of shallow cost gradient, high measurement noise, or low execution variability—the nervous system may choose to exploit its current control policy because whether it should adapt, and if so how it should adapt, is simply not clear.

**Figure 1:**
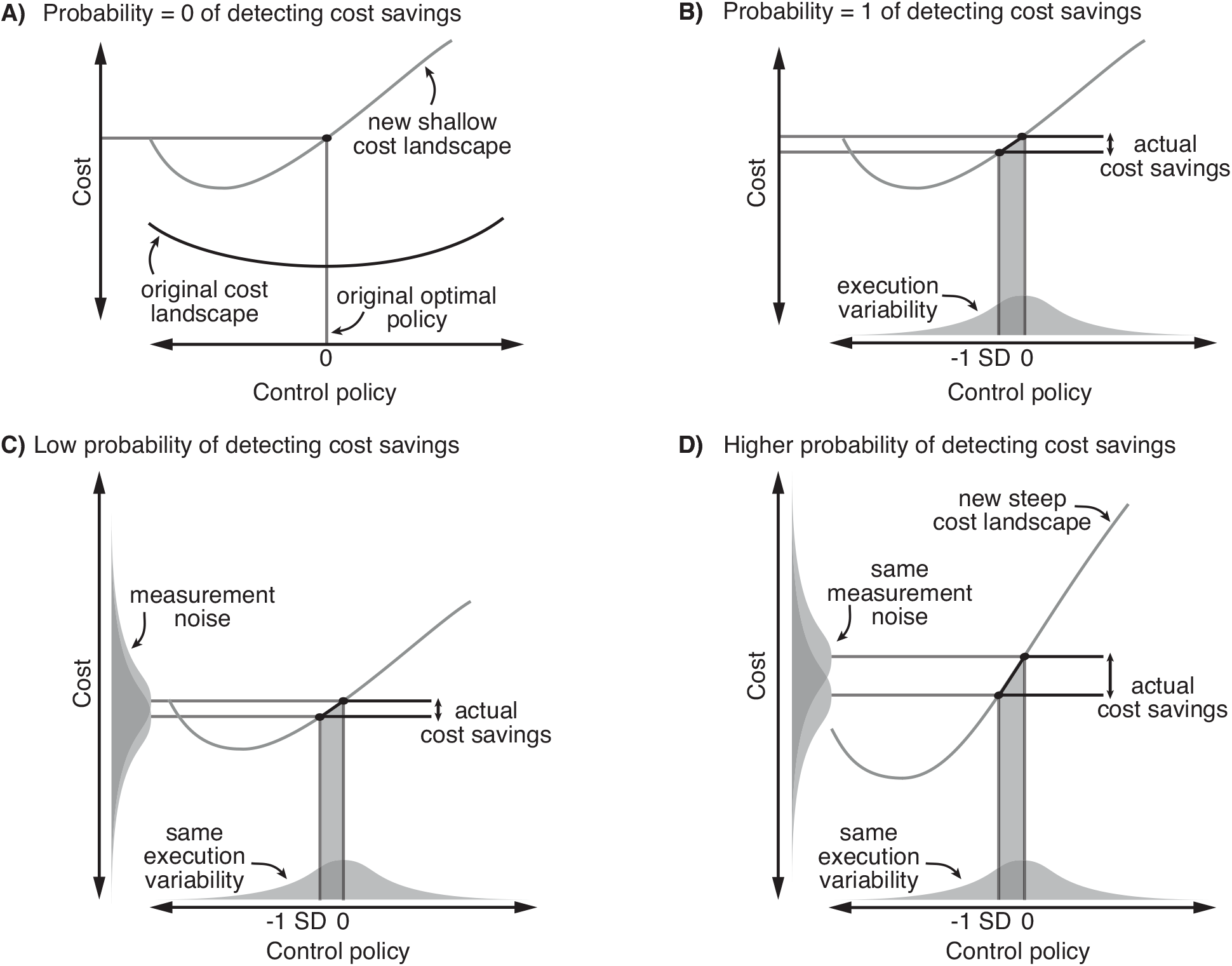
Conceptual representation of how the nervous system might detect cost savings from a cost landscape. **A)** The nervous system is introduced to a novel situation where the relationship between the control policy and cost has changed (black to grey curve) such that the original optimal policy is no longer optimal. With exact execution and measurement, the nervous system cannot detect any cost savings in the new landscape. **B)** Execution variability—illustrated by the horizontally aligned Gaussian distribution— allows the nervous system to exactly experience the lower costs relative to the original policy, making the energetic cost savings salient **C)** The presence of measurement noise—illustrated by the two vertically-aligned Gaussian distributions centered on the means of the two cost measurements—can reduce saliency by reducing the probability that the nervous system can detect a cost savings. In this example, the cost measurement means are close, and the cost measurement noise distributions are wide resulting in a low probability that the nervous system will detect a cost savings for the given execution variability. **D)** An increased gradient can increase the probability of detecting cost savings and thus increase the saliency of a cost landscape for the same execution variability and measurement noise.

Recent studies in walking support the premise that the nervous system relies on salient cost savings to initiate adaptation. One of the primary real-time objectives of the nervous system during walking is to minimize energetic cost (Abram et al., 2019; Selinger et al., 2019, 2015; Simha et al., 2019). In one of our recent studies, we used robotic exoskeletons to reshape the energetic cost landscape of treadmill walking. Here *cost landscape* refers to the relationship between step frequency and metabolic energetic cost. We reshaped the cost landscape to shift the optimal step frequency to step frequencies lower than normally preferred. Upon their first experience with the new cost landscape, only some participants spontaneously initiated adaptation to the new optimal step frequency. These *spontaneous initiators* had greater step frequency variability than the *non-spontaneous initiators* who persisted walking at the previous optimal step frequency. This suggests that the naturally higher variability increased the saliency of the cost savings to the nervous system which led to the initiation of adaptation. We were also able to prompt the non-spontaneous initiators to initiate adaptation by providing them with experience with step frequencies that resulted in a lower energetic cost. One interpretation of this result is that the experience increased the saliency of the energetic cost savings for the nervous system causing it to initiate further exploration. Counter to these findings, we did not find that increased gait variability was sufficient to initiate adaptation in a subsequent study on over ground walking (Wong et al., 2019). When compared to our treadmill studies, changes in cost in this over ground study were due not only to changes in step frequency, but also speed and terrain. We suspect that the nervous system did not initiate adaptation within the duration of this over ground experiment because the added dimensionality increased the complexity of the credit assignment problem making it difficult for the nervous system to determine which energetic changes could be attributed to its control, and which were due to the differences in terrain.

In the present study, we aimed to test whether the saliency of energetic cost savings is a feature that the nervous system uses to initiate adaptation in human walking. To accomplish this, rather than manipulate measurement noise or movement variability, we changed saliency by manipulating the gradient of the energetic cost landscape. We manipulated the gradient using a mechatronic system that applied controlled fore-aft forces to the waist of participants as they walked on a treadmill. These applied forces were a function of participants’ step frequency and acted to increase energetic cost at high step frequencies and reduce it at low step frequencies. By making the forces a function of only step frequency and keeping the walking speed constant, we aimed to only affect the gradient of the step frequency cost landscape, indirectly signaling to the nervous system how it should adapt its control policy to obtain cost savings. We increased the gradient of the cost landscape about participants’ originally preferred step frequency by increasing the magnitude of force change that the system provided for a given change in step frequency. We hypothesized that increasing the gradient of the cost landscape will cause participants to spontaneously initiate adaptation of their step frequency.

## Methods

### Experimental design

We manipulated cost landscapes using our recently developed mechatronic system (Figure 2A). We describe this system in detail in our earlier paper (Simha et al., 2019). Briefly, it manipulates a participant’s original cost landscape by applying fore-aft forces to their waist while they walk on a treadmill. The controller specifies the forces as a function of the participant’s step frequency. Backward forces increase the energetic cost associated with the executed step frequency, relative to normal, while moderate forward forces decrease the energetic cost (Gottschall and Kram, 2003). The system uses inertial measurement units placed on participants’ feet to detect ground contact events, and this signal is processed by a real-time controller to determine the participants’ executed *step frequency*, defined as the inverse of the time elapsed between left and right foot ground contact events. We provide the controller with a *control function* that defines the relationship it has to maintain between the measured step frequency and the applied force. Based on this control function and the measured step frequency, the controller commands the required force for each new step to an actuator via a motor driver. The force applied by the actuator is transmitted to the participants through long tensioned cables that are attached to a hip belt, and we monitor that force using force transducers in-line with the front and back cables.

**Figure 2:**
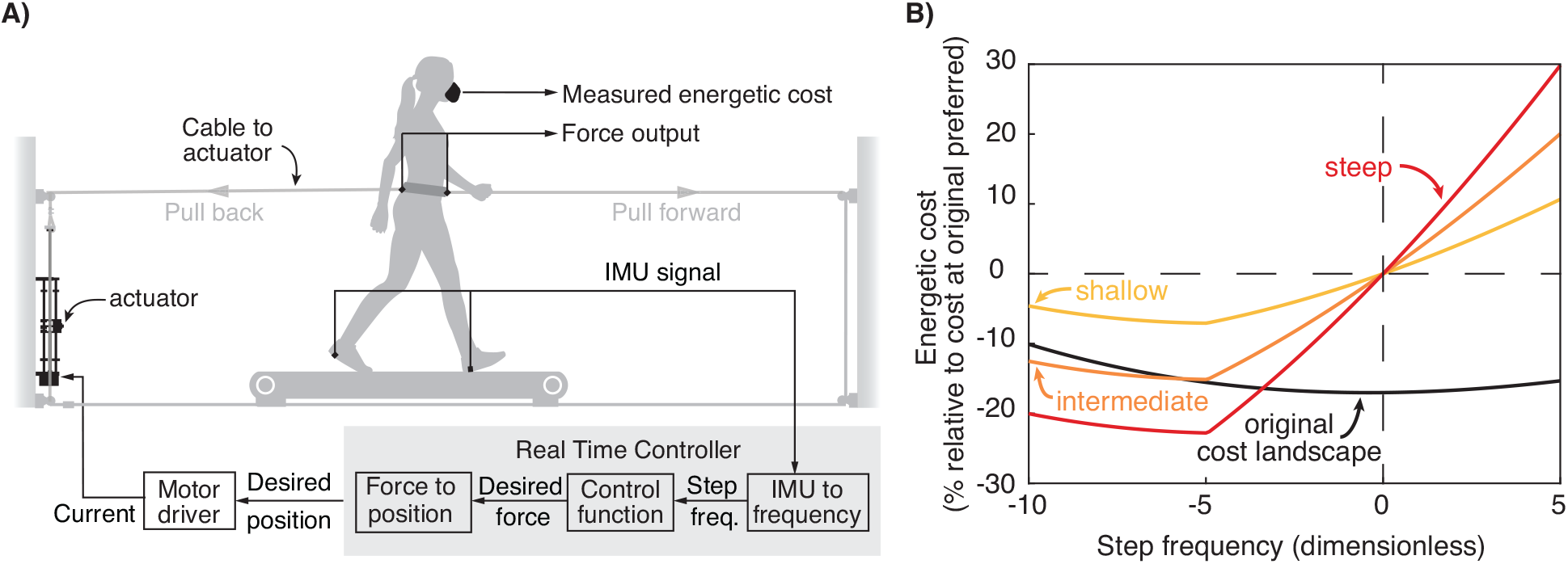
**A)** Participants walked in a mechatronic system that applied controlled fore-aft forces as a function of their walking step frequency. Backward forces provided an energetic penalty, raising the cost of walking relative to normal. Moderate forward forces provided an energetic reward, lowering energetic cost. **B)** Using simulations, we predicted that participants would experience cost landscapes with gradients of 1.4 (shallow), 2.8 (intermediate), and 4.2 (steep) percentage change in cost per unit change in step frequency, about their originally preferred step frequency (0).

We tested participants’ behavior in cost landscapes of three different gradients (Fig 2B). Using data from literature, we can predict for an average participant the energetic cost associated with each step frequency when walking without any external force (Umberger and Martin, 2007), as well as the energetic cost of walking when a range of fore-aft forces are applied but at a fixed step frequency (Gottschall and Kram, 2003). We combined these relationships and used them to design three control functions—*shallow, intermediate,* and *steep—*that created cost landscapes of three different gradients.

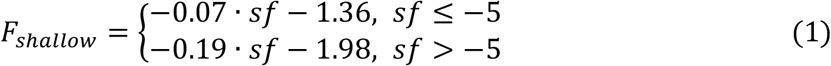

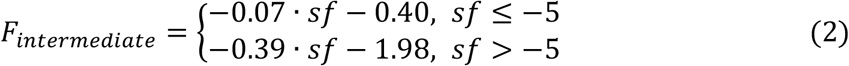

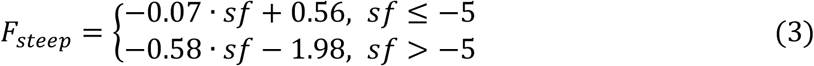

Here, *sf* is a normalized step frequency and is dimensionless. To perform this normalization, we first measured the average step frequency originally preferred by each participant during a baseline trial (c.f. Experimental Protocol), as well as the standard deviation in step frequency about this average preferred step frequency. We then calculated the normalized step frequency for each step in the subsequent trials by subtracting the average originally preferred step frequency from each step’s measured step frequency and then dividing by the standard deviation about the originally preferred step frequency. This normalization controls for the differences between participants in their step frequency variability, which is normally one of the contributors to the saliency of cost savings. It also forces measured step frequencies that are equal to the originally preferred step frequency to evaluate to 0. We normalized the forces applied to a participant by their body weight. In equations 1-3, the intercepts, slopes, and the forces all have units of percent body weight (*sf* is dimensionless). We designed these control functions to generate new cost landscapes with cost gradients of 1.4, 2.8, and 4.2 about the originally preferred step frequency. These gradients have units of percent change in energetic cost for a unit change in normalized step frequency. For example, were a participant walking in the intermediate gradient condition to choose a step frequency 1 normalized unit lower than their originally preferred step frequency, the participant will experience a 2.8% reduction in energetic cost relative to what they experienced at the originally preferred step frequency. For comparison, the cost landscape used in Selinger’s study roughly corresponds to the shallowest gradient we use here (Selinger et al., 2019, 2015). To experience a cost gradient as steep as our steepest, one would have to walk at a step frequency roughly 7.5% higher than their preferred step frequency in their original cost landscape (Umberger and Martin, 2007). Finally, we designed all the new cost landscapes to have the same cost at the originally preferred step frequency. This helped ensure that when we changed the cost landscape, participants only experienced the gradient change, without experiencing any change in the average steady-state cost. Since different nervous systems can respond differently to our control functions, each participant may not experience exactly the cost landscape that we aimed to create. However, our prior results show that, on average, we are able to accurately create our designed cost landscapes (Simha et al., 2019).

### Experimental protocol

We collected data from 11 participants (mean±SD; Age: 24±3 years; Height: 167±11 cm; Mass: 68±11 kg; Gender: 5 females, 6 males). All participants were healthy and had no known history of cardiopulmonary or gait impairments. The study protocol was approved by the Simon Fraser University Research Ethics Board and all participants gave written informed consent before participation. To determine the sample size necessary to evaluate our hypothesis, we first performed pilot experiments and estimated that we can expect a group standard deviation of 1.01 steps per minute. We then performed a power analysis for a one-tailed Students’ t-test to detect an average change of 1 normalized unit in step frequency (α = 0.05, 1-β = 0.90).

Each participant completed four periods of walking on the same day (Figure 3). Prior to the beginning of these four experimental periods, all participants spent ~10 minutes habituating to walking on our treadmill at a speed of 1.25 m·s^−1^. During this habituation, we instructed them to walk with both short and long steps. They were not attached to the mechatronic system. This was followed by the first period of the experiment where participants walked for 9 minutes while attached to the mechatronic system. We used data from this period to quantify the characteristics of their baseline walking step frequency. During this time, the system controlled for a target applied force of 0 N (Figure 3A). We calculated the average and standard deviation of their step frequency from the 6^th^ to 9^th^ minute to parameterize the step frequency in future trials. We refer to this average as the *originally preferred step frequency* and the standard deviation as *original step frequency variability*.

**Figure 3:**
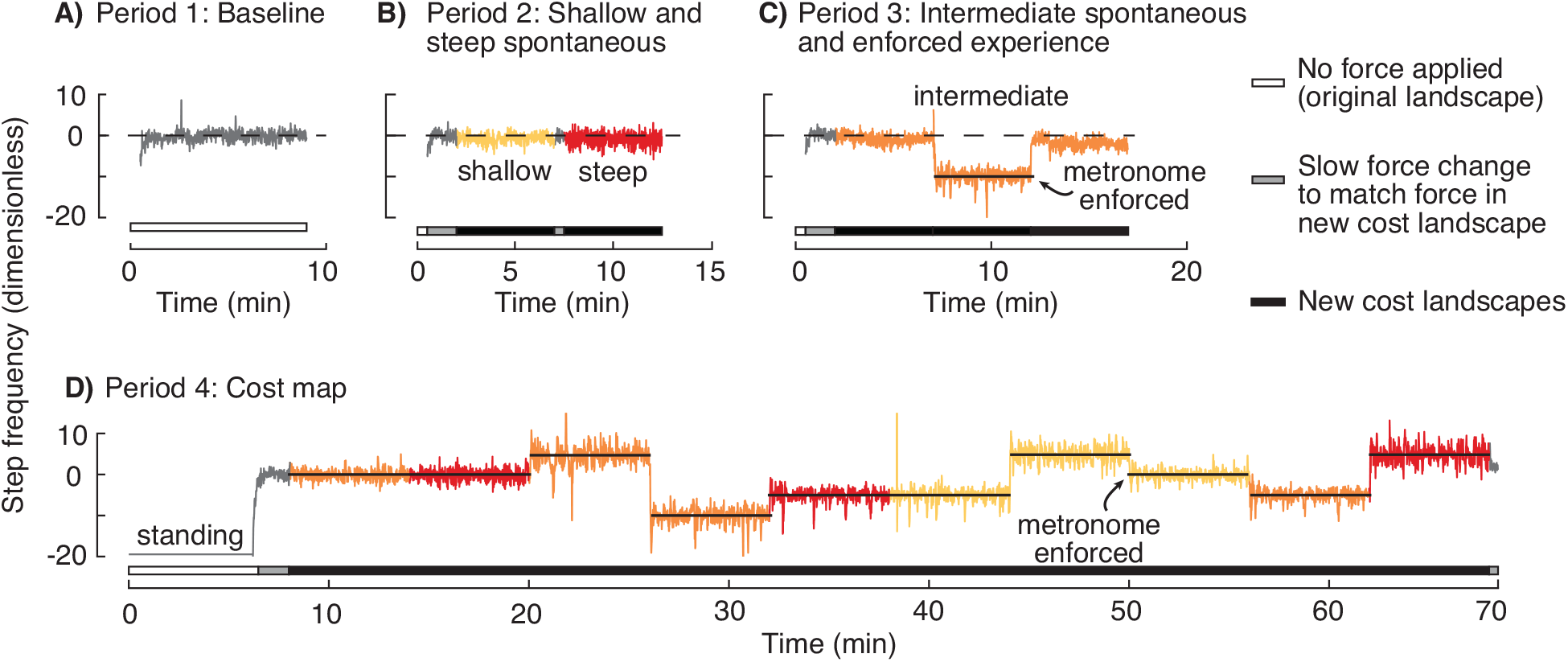
Step frequency measured from a representative participant during the different walking periods. Each participant completed four periods of walking in a single day. **A)** First, they walked for 9 minutes as the system controlled for a force of 0 N to be applied to their waist. We used this baseline period to estimate their average original preferred step frequency and original step frequency variability**. B)** Then participants walked for 5 minutes each in the shallow and steep gradients to test for spontaneous adaptation. **C)** In the third period, participants walked in an intermediate gradient. We used this condition to test for both spontaneous adaptation to an intermediate gradient and adaptation after enforced experience with a low cost. **D)** Finally, we measured the actual gradients experienced by participants in each the cost landscapes.

In the second period, we tested whether participants would spontaneously initiate adaptation in the shallow and steep gradients (Figure 3B). They experienced 0 N for the first 30 s to allow them to reach a steady-state step frequency (Pagliara et al., 2014). We programmed the system to ramp up the force over the next minute (minute 0.5 to 1.5) to the force that would be applied at the participants’ originally preferred step frequency in the new cost landscapes. This ensured that participants were not perturbed by a sudden change in force when the cost landscape changed. This force was held constant for 30 s (minute 1.5 to 2; *shallow pre-spontaneous*). The controller then engaged the control function for the shallow gradient, and participants walked at a self-selected step frequency for 5 minutes (minute 2 to 7; *shallow spontaneous*). Then the controller switched to the steep gradient. Once again, we ensured that participants were not perturbed during the cost landscape transition by using a limit on the rate at which the force could change for 30 s (minute 7 to 7.5). Participants then self-selected their step frequency for five minutes (minute 7.5 to 12.5; *steep spontaneous*). To avoid fatigue, we then provided a break of 5-10 minutes before beginning the third period. For each participant, we averaged their self-selected step frequency over the last 30 s of walking in each gradient to determine their *spontaneous adaptation* in that gradient (*shallow spontaneous*: minute 6.5 to 7; *steep spontaneous*: minute 12 to 12.5).

We used the third period to test for adaptation in an intermediate gradient (Figure 3C). The first part of this period served as a sort of Goldilocks test in the event that the shallow and steep gradients were both perceived as extreme by the nervous system (Kidd et al., 2012). Similar to the second period, the force was ramped up in the first two minutes to prevent perturbing forces. The controller then engaged the control function for the intermediate gradient, and participants self-selected their step frequency for 5 minutes (minute 2 to 7; *intermediate spontaneous*). One possible outcome of our experiment was that participants would not spontaneously adapt in any of the gradients. With this outcome, we would not be able to distinguish between the possibility that participants will adapt but not spontaneously, and the possibility that participants won’t adapt at all in our system with our experimental paradigm. Therefore, the next part of this experimental period was to verify whether adaptation was possible at all. Prior work has shown that experiencing a lower cost in a new cost landscape is sufficient to cause the nervous system to initiate adaptation (Selinger et al., 2019). Using this principle, we next required participants to match their step frequency to an audio metronome that played a frequency −10 normalized units away from their originally preferred step frequency. According to our designed cost landscape, we expected this step frequency to provide a cost savings of 12.5% relative to the cost at 0. After five minutes of matching the metronome (minute 7 to 12; *intermediate metronome guided*), the metronome was turned off and participants self-selected their step frequency for another five minutes (minute 12 to 17; *intermediate post-experience*). Once again, we averaged each participant’s step frequency during the last 30 s of each condition to determine their preferred step frequency in that condition (*intermediate spontaneous*: minute 6.5 to 7; *intermediate post-experience:* minute 16.5 to 17).

The purpose of the fourth period was to measure the actual energetic cost experienced by the participants in each of the new cost landscapes (*cost mapping*; Figure 3D). During this period, participants were also instrumented with a respiratory gas analysis system (Vmax Encore Metabolic Cart, Viasys, Pennsylvania, USA). They spent the first six minutes standing still while we measured their resting metabolic rate (minute 0 to 6). They then started walking while the mechatronic system maintained a force of 0 N to allow them to reach a steady-state gait (minute 6 to 7). Following this, participants walked at specific walking conditions chosen to allow us to estimate the gradient about the originally preferred step frequency in each of the cost landscapes, and also to estimate if the *experience low* period indeed allowed participants to experience a lower cost. Participants walked in 10 conditions total: step frequencies of 0, −5, and +5 in shallow, intermediate and steep gradients, and −10 in only the intermediate gradient. We enforced this by instructing participants to match an audio metronome that played these frequencies. We programmed the controller to present these conditions in a random order to each participant, to prevent any order effects on these metabolic energy measures. To determine energetic cost, we measured the total volume of oxygen consumed and volume of carbon dioxide produced in the last three minutes of each condition, and divided them by the duration over which they were measured, to obtain the steady state average rates of oxygen consumption 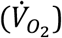 and carbon dioxide production 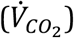. We then estimated the metabolic rate using the following equation (Adamczyk et al., 2006; Brockway, 1987; Weir, 1949):

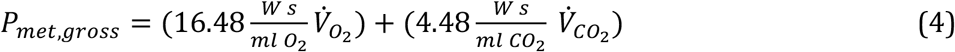

We subtracted resting metabolic power for each participant and present net energetic cost as the energy used per unit time normalized for the person’s body mass. It has the units W·kg^−1^.

### Data analysis

We first determined the average gradients and metronome-guided cost that participants experienced in each cost landscape. We used MATLAB’s fitlm command to find the best linear fit through the energetic costs at −5, 0, +5 normalized step frequencies for each participant, in each cost landscape. We define the cost landscape gradient for each participant as the slope of this fit. We also used a one-tailed paired Students’ t-test to determine whether the cost at a step frequency of −10, where we held participants during the *intermediate metronome guided* condition, is lower than the cost at a step frequency of 0 in the intermediate gradient.

We evaluated whether participants spontaneously initiated adaptation in response to steeper gradients. We first compared the preferred step frequency in the shallow gradient with the originally preferred step frequency using a one-tailed Student’s t-test. We found that these values were indeed different, but we did not attribute this shift in preferred step frequency to an adaptation in response to a new cost gradient (c.f. Results). To determine if there was any additional changes in preferred step frequency in the steeper gradients, we then compared the average step frequencies during the spontaneous adaptation periods in the intermediate and steep gradient to the same period in the shallow gradient.

We also determined whether participants initiated adaptation after enforced experience with a low cost. We used a one-tailed paired Students t-test to determine whether participants’ preferred step frequency after the *intermediate metronome guided* condition was significantly lower than the average step frequency 30 s prior to the experience with low cost. The step frequency in this 30s prior to the experience corresponds to the spontaneous adaptation in the intermediate gradient, allowing us to determine whether the metronome guided experience generated adaptation that did not occur spontaneously.

In the conditions where we observed adaptation, we characterized the rate of adaptation. We did so because a preferred step frequency can arise from fast predictive processes that can occur over a few seconds or optimization processes that can occur over tens or hundreds of seconds (Pagliara et al., 2014). As described in the introduction, we are interested in the slow process since it is indicative of the nervous system learning to adapt its policy to a novel situation. We modelled each participant’s adaptation of step frequency over time as a two-process exponential. We first averaged the step frequency during the last 30 s prior to the beginning of the condition of interest, and the step frequency during the last 30 s of the condition. If these two averages were different, we normalized the step frequency data during that condition such that the average step frequency of the 30 s prior to the condition evaluated to 0, and the average of the last 30 s evaluated to 1. We then used least squares regression implemented through MATLAB’s fitnlm function to model these data as the sum of two exponentials (Pagliara et al., 2014). We used the time constants from this model to estimate the duration of the optimization process.

## Results

We were successful in creating cost landscapes of different gradients. We found that participants on average experienced a shallow gradient of 0.07±0.03 W·kg^−1^ (mean±SD), an intermediate gradient of 0.14±0.03 W·kg^−1^, and a steep gradient of 0.20±0.04 W·kg^−1^ (Figure 4). This is calculated as the change in energetic cost per normalized unit of step frequency. We use 1 standard deviation of participants’ preferred step frequency in their original cost landscape, to normalize the measured step frequency. This means that participants experience the reported gradient through a variability of 0.5 standard deviations higher and lower than their originally preferred step frequency. Thus, 1 standard deviation higher and lower than their originally preferred step frequency, which accounts for 68% of their steps, would have allowed participants to experience a change in energetic cost of 3.6%, 7.2%, and 10.2% in the shallow, intermediate, and steep gradients, respectively. We also found that when participants in the intermediate gradient were held - 10 normalized step frequencies lower than their originally preferred step frequency, they experienced an average cost savings of 8.1%±9.1% relative to the cost at the originally preferred step frequency (p=0.006).

**Figure 4:**
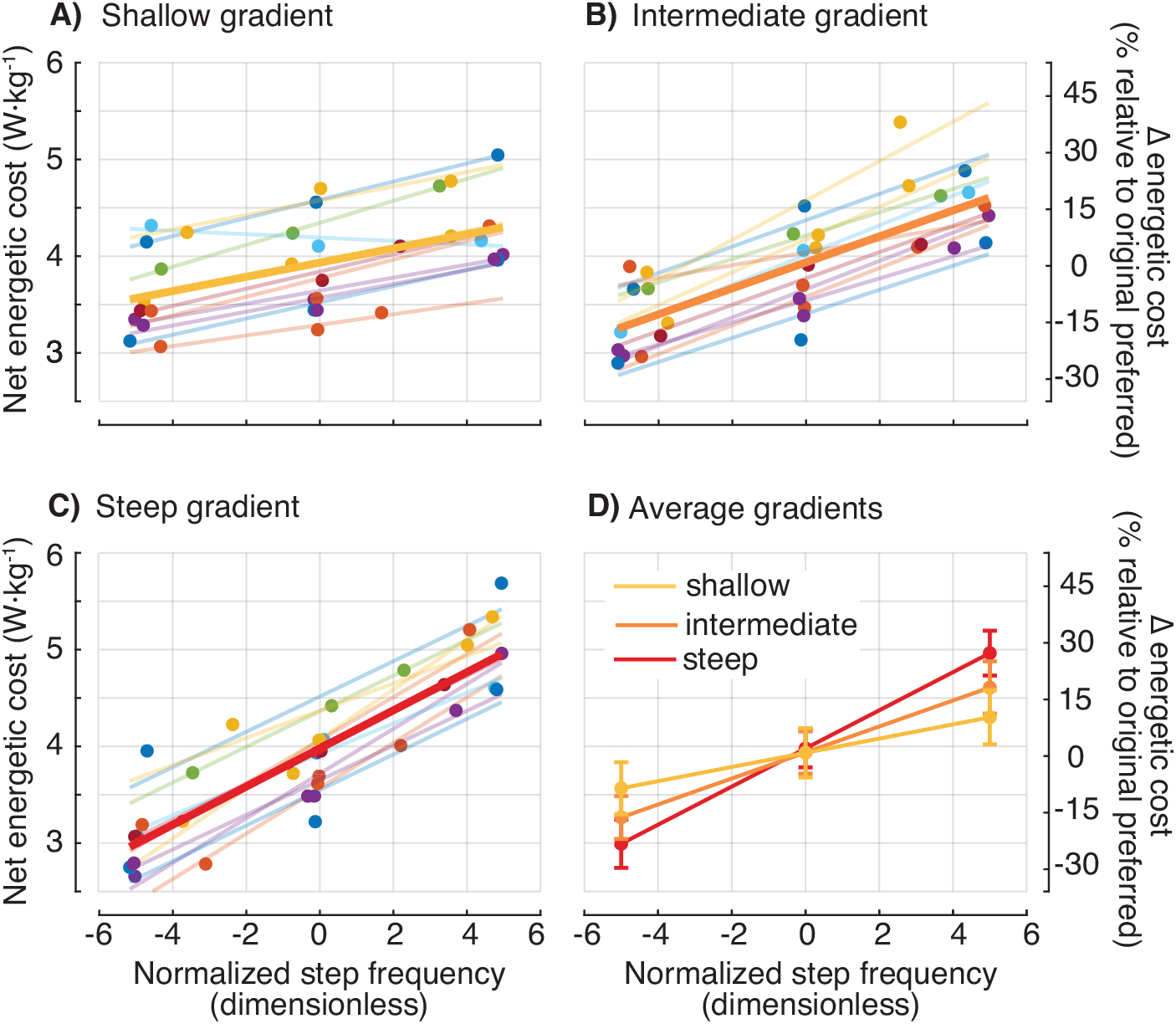
**A)** Shallow **B)** intermediate and **C)** steep gradients. Each filled circle represents one measurement from one participant. Light lines are linear fits to each participant’s cost measurements. Data points and best-fit lines from a given participant is presented in a single colour. Thick lines are the average of these linear fits. **D)** On average the gradients are increasing from shallow to steep. The filled circles represent the average cost measures at the commanded step frequencies, and the error bars represent the 95% CI of the same.

Participants did not spontaneously initiate adaptation in response to steeper cost gradients. In the second period, participants first experienced the shallow cost landscape, and then the steep cost landscape. They walked freely at their self-selected step frequency for 5 min in both cost landscapes. The average step frequency from the last 30 seconds of the shallow period was lower than the original preferred step frequency (−0.69±0.82 vs. 0; p = 0.01). However, this step frequency was indistinguishable from the average step frequency preferred by participants during the 30 s prior to the beginning of the shallow gradient (Figure 5: Shallow pre-spontaneous vs Shallow spontaneous; −0.69±0.82 vs. - 1.02±0.64; p = 0.33). Therefore, we do not interpret this to be an initiation of adaptation towards the optimal policy. When the system switched from the shallow landscape to the steep landscape, participants still did not initiate adaptation, and preferred a step frequency (Figure 5: Steep spontaneous; −0.76±0.99) that was indistinguishable from that preferred in the shallow cost landscape (Figure 5: Shallow spontaneous; −0.69±0.82; p = 0.40). Our goldilocks test with the intermediate gradient also resulted in preferred step frequencies that were indistinguishable from that preferred in the shallow landscape (Figure 5: Shallow spontaneous vs Intermediate spontaneous; −0.69±0.82 vs. −0.74±0.82; p = 0.43).

**Figure 5:**
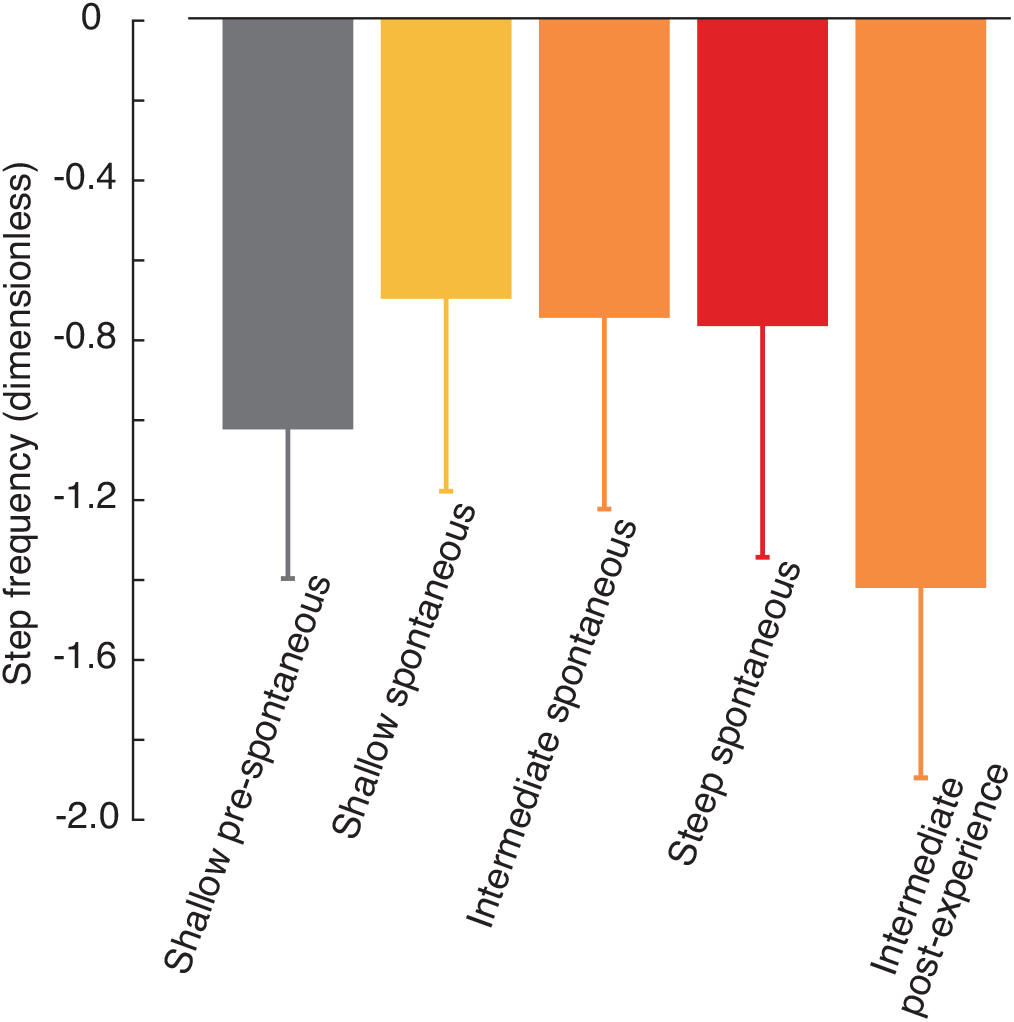
Average spontaneous adaptation in the shallow gradient was indistinguishable from participants’ preferred step frequency just prior to the beginning of the shallow cost landscape (shallow pre-spontaneous vs shallow spontaneous). The spontaneous adaptation in all gradients were also indistinguishable from each other after five minutes of walking (shallow, intermediate, steep spontaneous). However, after experience with a lower cost in the intermediate gradient, participants preferred to walk at a significantly lower step frequency (intermediate post-experience).

Participants did, however, initiate adaptation after enforced experience with a lower cost. We allowed participants to self-select their step frequency after matching a metronome that held them at a step frequency that had a cost lower than the cost at 0 in the intermediate cost landscape. On average, participants adapted by −1.41±0.81 towards the new cost minimum (Figure 5: Intermediate post-experience). This adaptation was to step frequencies significantly lower than that spontaneously preferred in the intermediate gradient (p=0.007). It led to an average cost savings of 4.80±3.12% relative to the cost at 0. We found that the time course of the change in step frequency of most participants was captured well with a two-process exponential model (RMSE=0.16±0.08; R-squared=0.36±0.21). The time constant of the fast process was 4.4±2.5 s while that of the slow process was 190.2±209 s. We interpret the presence of this slow process as evidence that the nervous system indeed initiated adaptation in response to the enforced experience with a lower cost gait.

## Discussion

Contrary to our hypothesis, steeper gradients did not lead to spontaneous initiation of adaptation. This null finding is not because our methods were unsuccessful in creating gradients of increasing steepness. We used our cost mapping trials to verify that the participants did indeed experience three different gradients—the intermediate and steep gradients were about 2-fold and 3-fold the shallow gradient, respectively. The lack of initiation also does not appear to be a consequence of the rapid exposure to multiple gradient conditions preventing the nervous system from attempting any adaptation. We verified this by leveraging results from previous studies that found that adaptation can be initiated by guiding the nervous system to experience a cost lower than the cost at the originally preferred step frequency (Selinger et al., 2019). We did the same here and found that participants could indeed initiate adaptation in the intermediate landscape after such experience, despite the intermediate landscape being the third landscape experienced by participants. When considered together, these results suggest that either the nervous system does not use salient cost savings to initiate adaptation, or that the cost savings were not salient to the nervous system in our experiment.

For savings to be salient, the nervous system needs to both detect that cost savings can be gained and determine how it should adapt its control policy to gain the savings. Depending upon how the nervous system senses energetic cost, it may be challenging for the nervous system to detect cost savings from the cost landscape gradient. For example, one possible candidate sensory system for estimating energetic cost involves the ergoreceptors that are sensitive to the slow build-up, or slow reduction, of muscle metabolic byproducts (Amann et al., 2011; Iwamoto et al., 1985; Mitchell et al., 1983). This build-up creates a sensory response that is an integration of the effect of many steps, rather than one that closely follows the step-to-step changes in energetic cost. It will be more difficult for the nervous system to detect a gradient in cost landscape from the step-to-step variability in energetic cost when using this mechanism because integration has the effect of decreasing the sensed gradient, perhaps even to zero if the build-up is particularly slow. This, or a similar integrative sensing mechanism, may be why metronome-guided experience is effective at initiating adaptation—the metronome holds participants at a lower cost for many steps allowing time for integration. However, some participants in some conditions are able to use the step-to-step variability in energetic cost to spontaneously initiate adaptation (Selinger et al., 2019). This suggests that if a slow sensing system does indeed play a role in estimating energetic cost, it is not the only contributing system.

Another possibility for the lack of initiation of adaptation is that the gradients allowed participants’ nervous systems to sense the presence of cost savings but not how to adapt their control policy to obtain those savings. That is, the nervous system has difficulty with credit assignment in our experiment (Guerguiev et al., 2020). We manipulated the cost gradient associated with only one gait parameter—step frequency—to allow the nervous system to detect an increase in cost savings and detect the gait parameter to adapt to obtain those savings. But when walking in our system, we suspect that it is not clear to most participants that the backward force depends on any aspect of their gait, including their step frequency. It appears to be challenging for the nervous system to identify salient cost savings using the structure of natural variability in gait to determine the gradient of a cost landscape—a finding consistent with our earlier experiment studying adaptation in over ground walking (Wong et al., 2019). Metronome-guided experience of step frequencies with lower cost may provide the nervous system with an explicit association between the cost savings and the changes to control policy that provide those cost savings. Similarly, reaching experiments have found that presenting participants with multiple different force-fields interferes with learning, but that such interference can be overcome with certain contextual cues such as follow through movements or cues that associate a change in the optimal control policy with another change such as spatial location of movement (Howard et al., 2015, 2013). Differences in contextual clues might explain why it was easier for the nervous system to identify that there was a relationship between step frequency and the changes to knee torque for some participants in our previous experiment than with step frequency and torso forces in the present experiment (Selinger et al., 2019). This interpretation is consistent with recent study in visuomotor adaptation that found that implicit and explicit learning work together to improve adaptation (Miyamoto et al., 2020).

While we designed our custom-built equipment and our protocol to meet the requirement for energetic cost saliency, our experiment nevertheless had limitations. Towards this requirement, the maximum cost savings that participants experienced from their variability in step frequency, relative to the cost at their originally preferred step frequency, was 5.1% in the steep gradient. In contrast, participants experienced cost savings of 8.2% during their metronome-guided lower cost experience in the intermediate gradient. This suggests that even the steep gradient may not have allowed participants to experience a large enough cost savings. However, we suspect this is not the case because in our previous study with the shallow gradient, participants initiated adaptation after experiencing cost savings of only 3.5% through similar metronome-guided walking (Simha et al., 2019). This earlier cost savings was smaller than that experienced by our current participants in the steep gradient condition suggesting that the currently experienced cost savings, at least in the steep gradient condition, were sufficiently large for the nervous system to detect.

A second limitation is that our experimental design resulted in participants preferring step frequencies slightly lower than the original preferred step frequencies in all gradient conditions (Figure 5). We do not interpret these shifts as evidence of the initiation of energetic cost optimization in response to new cost landscapes. Our rationale is that participants were already walking at shifted step frequency during the 30 s prior to the beginning of each new cost landscape (shallow: −1.02±0.64, intermediate: −0.74±0.82, steep: −0.62±0.54). Why is step frequency shifted lower than the baseline measures both before and during the experience with the new cost landscapes? One possible explanation is that we may not have provided a long enough baseline period for participants to settle into their preferred step frequency. However, others have found that two minutes of walking is sufficient for stride frequency to reach steady state—we provided 9 minutes (Van de Putte et al., 2006; Wall and Charteris, 1981). A second possible explanation for the presence of these shifts may be the net backward force that participants experienced both immediately before and during the cost landscape, but not during the baseline phase when the net force was zero. Our system slowly ramped up the backward force to that which participants would experience in the new cost landscapes at their originally preferred step frequency. The force was then held constant for 30 s before the controller switched to the new cost landscapes, and our step frequency estimate prior to the beginning of the new cost landscape is from this constant-force period. However, concerned about the possible role of net backward force on step frequency, we performed pilot experiments prior to our reported experiments and found no relationship. In support of our pilot results, a recent study also found that backward forces do not have an effect on stride period (Dewolf et al., 2020). Furthermore, walking uphill, which is biomechanically similar to experiencing a net backward force, also results in step frequencies that are not significantly different from walking on level (Ortega and Farley, 2015). Further research will be required to understand why we observed this consistent shift in step frequency.

After metronome-guided experience, our participants did not converge on the energy minimal step frequency. To determine the location of the cost minimum, and the magnitude of cost savings obtainable at the minimum, we fit a quadratic relationship to the costs measured in the intermediate gradient condition during cost mapping. From this relationship, we estimate that, on average, participants could have obtained a cost savings of 10.8% if they had shifted their step frequency −6.1 normalized units away from their originally preferred step frequency. Yet we found that participants only adapted their step frequency by −1.4±0.8 to obtain a cost savings of only 4.83±3.61%. This might suggest to some that energetic cost savings do not play a role in the adaptation of step frequency in the cost landscapes we used. We suspect that this is not the case since participants did adapt after experience with a lower energetic cost. A candidate explanation is that the nervous system seeks to minimize an objective function that is a combination of energetic cost, stability, accuracy and other contributors (Abram et al., 2019). The minimum of this combined cost function may coincide with the final preferred step frequency and not with the energetic cost minimum.

In conclusion, the nervous system does not solely rely on the gradient of energetic cost to initiate adaptation in novel situations. As we and others have previously found, explicit experience with more optimal movements can assist with the initiation of adaptation. A better understanding of the interplay between implicit and explicit experience for the nervous system to initiate adaptation when the saliency of cost savings is not apparent may help improve rehabilitation for those recovering from injuries, help coaches speed up training with new techniques, or aid scientists looking to study adaptation in complex novel environments.

